# The Global Wheat Full Semantic Organ Segmentation (GWFSS) Dataset

**DOI:** 10.1101/2025.03.18.642594

**Authors:** Zijian Wang, Radek Zenkl, Latifa Greche, Benoit De Solan, Lucas Bernigaud Samatan, Safaa Ouahid, Andrea Visioni, Carlos A. Robles-Zazueta, Francisco Pinto, Ivan Perez-Olivera, Matthew P. Reynolds, Chen Zhu, Shouyang Liu, Marie-Pia D’argaignon, Raul Lopez-Lozano, Marie Weiss, Afef Marzougui, Lukas Roth, Śebastien Dandrifosse, Alexis Carlier, Benjamin Dumont, Benôıt Mercatoris, Javier Fernandez, Scott Chapman, Keyhan Najafian, Ian Stavness, Haozhou Wang, Wei Guo, Nicolas Virlet, Malcolm J Hawkesford, Zhi Chen, Etienne David, Joss Gillet, Kamran Irfan, Alexis Comar, Andreas Hund

## Abstract

Computer vision is increasingly used in farmers’ fields and agricultural experiments to quantify important traits. Imaging setups with a sub-millimetre ground sampling distance enable the detection and tracking of plant features, including size, shape, and colour. Although today’s AI-driven foundation models segment almost any object in an image, they still fail for complex plant canopies. To improve model performance, the global wheat dataset consortium assembled a diverse set of images from experiments around the globe. After the head detection dataset (GWHD), the new dataset targets a full semantic segmentation (GWFSS) of wheat organs (leaves, stems and spikes) covering all developmental stages. Images were collected by 11 institutions using a wide range of imaging setups. Two datasets are provided: i) a set of 1096 diverse images in which all organs were labelled at the pixel level, and (ii) a dataset of 52,078 images without annotations available for additional training. The labelled set was used to train segmentation models based on DeepLabV3Plus and Segformer. Our Segformer model performed slightly better than DeepLabV3Plus with a mIOU for leaves and spikes of ca. 90%. However, the precision for stems with 54% was rather lower. The major advantages over published models are: i) the exclusion of weeds from the wheat canopy, ii) the detection of all wheat features including necrotic and senescent tissues and its separation from crop residues. This facilitates further development in classifying healthy vs. unhealthy tissue to address the increasing need for accurate quantification of senescence and diseases in wheat canopies.

## 1 Introduction

Wheat is one of the most important crops in the world, providing 18% of calorie intake and 19% of protein intake globally [1]. While the area of farmland used to grow wheat has remained stable, global wheat yields have quadrupled since 1960, mainly due to technical innovation, such as the widespread use of N fertilisers, and the breeding of modern wheat varieties [1]. However, the rate of yield gain has stagnated or even decreased in the last two decades in different regions of the world [2], imposing a challenge to fulfilling the projected demands for wheat production in the future [3]. In addition, climate change-induced stresses [1], plant disease adaptation, and pest outbreaks, combined with the need for more efficient crops that require less input in terms of fertilisers, water, and pesticides place additional constraints on wheat production. Increasing wheat yield is a multi-faceted problem which involves genetic, physiologic, and agronomic improvement to enhance resource-use efficiency. Novel phenotyping approaches provide advanced tools and methodologies to enhance wheat management, optimise breeding, and achieve efficient resource utilisation. These methodologies are key to enabling precise repeatable measurements in agricultural fields and research networks across the globe, as highlighted in a survey on field-based phenotyping in Europe [4]. In their survey, Morisse et al. [4] emphasise the capability of these platforms to enable the rapid collection of datasets at breeding scale, including hundreds of plots with various genotypes and treatments. Moreover, data can be collected during the entire crop growth cycle, enabling tracking of wheat responses to biotic and abiotic stresses or management practices. Although a wide range of possible sensors are available, including 3D scanners [5] to characterise canopy architecture or hyperspectral imaging [6, 7] to monitor plant health and productivity, we focus here on high spatial-resolution imaging with red, green and blue (RGB) spectra. Such imaging provides a broad range of phenotyping capabilities at relatively low equipment costs and high spatial resolution. In a comprehensive review on translating high-throughput phenotyping (HTP) into genetic gain, Araus et al. [8], posed a key question: *“Will low-cost HTP tools be adopted regularly by breeders in the next decades? If so, are RGB cameras, mobile apps, and drones the natural candidates?”* Classical RGB cameras can be mounted on handheld devices [9], ground-based vehicles [10] [8], gantries [11], or drones [12], making them adaptable to various scales platforms. The advantage of RGB sensors is their relatively high spatial resolution and low costs compared to other HTP tools [13]. This enables the capture of sufficient details to separate plant features from complex canopies. However, robust feature extraction requires algorithms capable of extracting information from images taken under a wide range of conditions. The development of such algorithms demands a sufficient number of well-annotated training images capturing this diversity.

Several pioneer datasets published related wheat-linked tasks in field conditions focusing either on (i) spike detection and quantification with the SPIKE [14] and GHWD dataset [15, 16], and a dataset provided by Madec et al. [17], (ii) vegetation segmentation on wheat only with EWS dataset [18] or on multispecies including wheat with SegVeg and VegAnn [9, 19] or (iii) disease detection on wheat with the NWRD dataset [20], the Wheat Leaf dataset for strip rust and septoria [21], the CDTS for strip rust [22], the Wheat nitrogen deficiency and leaf rust image dataset [23] and the CGIAR Computer Vision for Crop Disease for stem and leaf rust [24]. Despite these contributions, existing datasets are often limited in terms of geographic diversity, genotype variations, and growth stages, which may limit the generalisation power of models trained on these datasets. Furthermore, only a few datasets are collaborative, involving multiple countries and institutions [9, 15, 16, 19]. To address this limitation, the collaborative Global Wheat Dataset Consortium was established. This consortium aims to aggregate datasets from multiple institutions, make them publicly available, and provide the necessary data to develop robust algorithms. In previous GWHD editions 2020-2021 [15, 16], we released a large dataset with a total of 6,515 high-resolution RGB images, containing annotations for 275,187 labelled wheat heads. The early design of GWHD focused on providing bounding-box annotations for wheat heads, including images of different genotypes captured under varying environmental, management and growth conditions. GWHD played a pivotal role in various research studies, particularly in developing and benchmarking wheat head detection and counting methods using supervised models [25–31], semi-self-supervised models [32, 33] and self-supervised models [34], generating a reference dataset to improve deep learning model performance [35–37] and improving head count models in dense plots [38]. These robust detection models allow the counting of wheat heads per unit area [29, 39], as long as the footprint of the image at the top of the canopy is known. In addition, the dynamics of head counts may be used to approximate heading dates [40]. The flowering date of wheat is often approximated by the heading date because it is easier to assess head emergence or presence than anther extrusion which is more affected by time of day, wind conditions and operator experience [40, 41]. Thus, heading is the most widely assessed trait related to cereal phenology and an important trait to understand the effect of environmental stresses, such as heat and drought, on grain yield [42].

While counting wheat heads is important as it is one of the yield components, it is not the only targeted trait. Wheat continuously adjusts its yield potential during the entire vegetation period. Low germination or plant damage due to winter kill is compensated for by increased tillering, while during stem elongation, excessive tillering is compensated by tiller abortion. Later in the season, the different organs can undergo different senescence dynamics, as demonstrated by Anderegg et al. [43]. A deeper understanding of how yield is formed throughout the growth season will benefit from a non-destructive assessment of its components. The imaging and semantic segmentation of all the plant organs visible in the image, from emergence to maturity, have great potential to shed light on the yield formation process. Examples of targeted traits are seedling count [44, 45], canopy cover [18, 19, 46, 47], biomass estimation [48] and leaf area index (LAI) [49, 50]. In most cases, segmentation of wheat canopies from other background features, such as soil or weeds, is required. The comparably simple task of segmenting canopies from the soil background was previously solved by manual adjustments [51], using automatic threshold methods, such as the Otsu algorithm [52, 53] or shallow machine learning [13, 54, 55].

The above-mentioned approaches based on colour information at the pixel-level, have the short-coming that they cannot take context into account. Modern deep learning methodologies enable the learning of contextual information. This requires human-annotated training data to supervise feature detection in complex images containing many plants growing together in a canopy. The different plant organs in such canopies are not simply green but might have different shades of green or yellow due to senescence, chlorosis, or necrotic tissues. Necrotic plant tissues may have a similar brown colour as crop residues and can only be segmented in RGB images based on context. Green canopy segmentation using deep learning models has become a standard procedure, with training data sourced from a wide range of crops, such as the VegAnn dataset [9] combining 3775 RGB images of 12 different crops. With the existence of large datasets and the advancements in computer vision, new possibilities of data processing and feature extraction have been unlocked through the utilization of data-driven deep learning approaches. Existing deep learning-based segmentation is mainly based on encoder-decoder architecture like DeeplabV3Plus [56–60] and Atrous convolution [61, 62] architecture, with application-specific adaptation. More recently, transformer-based segmentation models (e.g., SegFormer [63]) have gained attention due to their ability to capture long-range dependencies and global context effectively. These models show promise in addressing complex segmentation tasks, offering improved performance and adaptability in agricultural applications. Despite their potential to achieve promising performance, transformer-based models require relatively larger datasets that are currently lacking in agricultural domains [64].

Within the wheat canopy, segmentation of organs including leaves, stems, and heads is required, followed by the extraction of relevant phenotypes from each organ. Many wheat researchers have collected their own wheat datasets or made their own annotations from existing ones to achieve semantic segmentation of heads [35, 59, 65–67], spikelets [68], grains [69], stems and foliage [43] infected [20, 70–72] and senescent [19] tissues or a combination of disease and senescence [73]. The data provided in these publications largely advance wheat phenotyping at the organ level, offering tools for detailed studies of yield components such as spike number, spikelets per spike, spikelet size, leaf disease resistance, and senescence dynamics. However, there is a lack of integrated datasets enabling simultaneous segmentation of all wheat organs (leaves, spikes, stems) from soil, crop residues, weeds and other background elements. Moreover, wheat has a complex canopy due to its high planting density, strong development of tillers (lateral shoots), thin stems, overlapping leaves and occluded organs. Variations in appearance caused by growth stages, lighting conditions, wind patterns and imaging angles make it a challenging plant species to phenotype. To advance our capabilities beyond the wheat head-centred GWHD dataset 2021-2022, we assembled the Global Wheat Full Semantic Segmentation (GWFSS) dataset collected by different phenotyping platforms from 11 institutes and universities across the globe under various light and weather conditions. This diversity ensures that the dataset addresses the extensive requirements of wheat phenotyping across a range of genetic backgrounds, environments, and management practices throughout the growing season. The images were collected with an average ground sampling distance (GSD) between 0.09 and 0.71 mm per pixel. This is significant for accurately capturing organ features to allow precise differentiation and measurement of these smaller structures, rather than focusing solely on canopy-level traits. Our contribution can be summarised as follows:

1. A *full* GWFSS dataset comprises 52,078 RGB images without labels is available in ETH research collection.
2. An *annotated* GWFSS dataset providing 1,096 pixel-level annotations (masks) for the following classes: leaves, stems, heads, and background.
3. The results of two state-of-the-art segmentation models, DeepLabV3Plus and Segformer, finetuned on the full dataset. The models were trained as a baseline performance benchmark for organ segmentation.

The 1,096 images are held-out test data for a data challenge, which will not be released with this pre-print. The data will be fully released on 2025-06-25, after the MLCAS25 competition has closed.

## 2 Material and Methods

### 2.1 Field Experiments

The dataset includes images from field experiments conducted by 11 institutions worldwide. Wheat plots at 67 different field sites were imaged using proximal RGB imaging setups throughout the growing seasons (Fig. 1). The experiments cover a wide range of planting densities, agronomic inputs, environmental conditions, as well as disease and weed pressures. Thus, the GWFSS dataset spans diverse agroclimatic zones and management practices. The imaging setups used by the 11 institutions and the data sets derived are described in detail in Table S1 and Table 2, respectively. Additional information related to the datasets is given as follows with institutions listed in alphabetic order:

**Figure 1:**
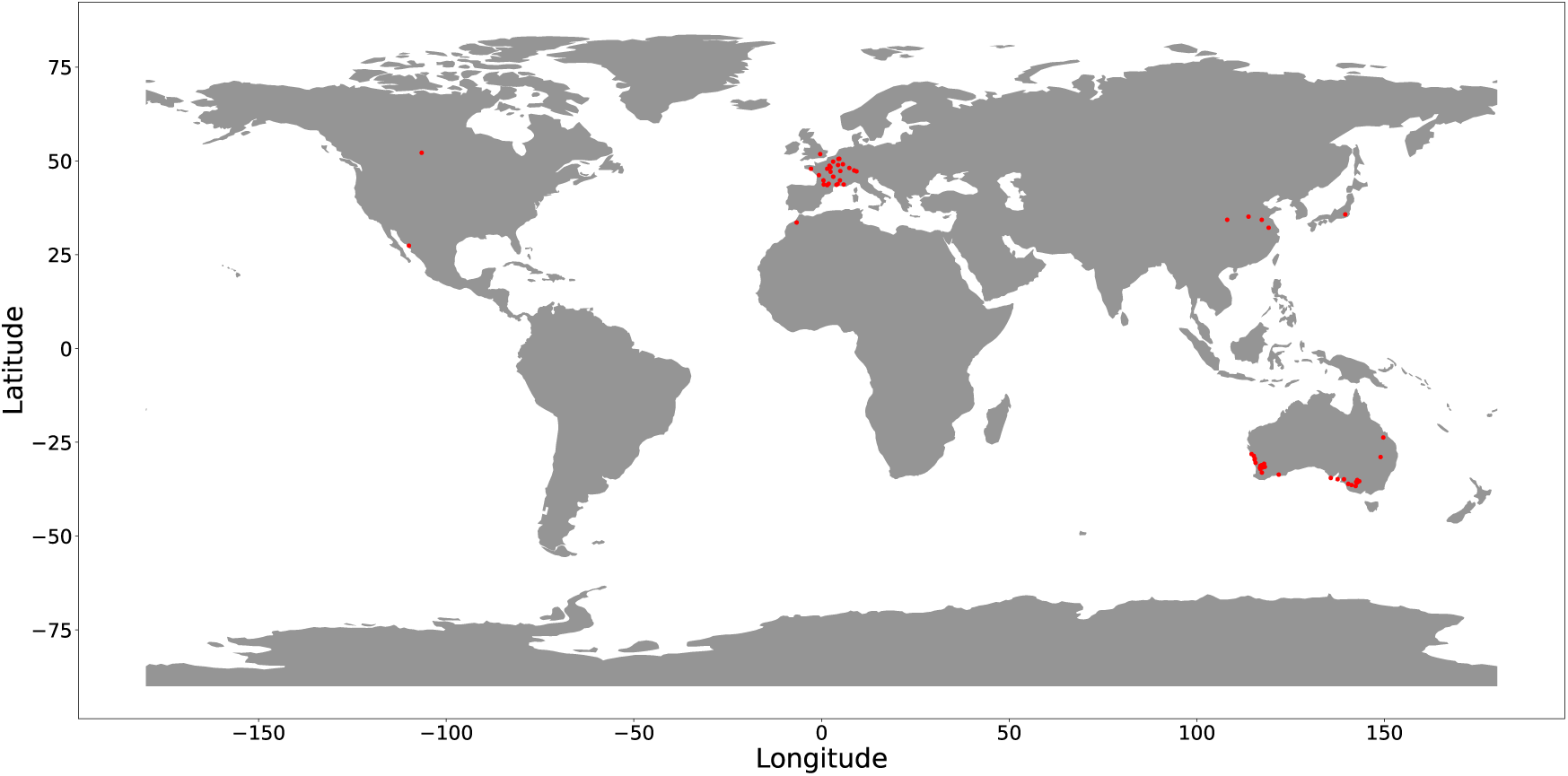
An overview of the location of all trials included in GWFSS.

**Table 1:**
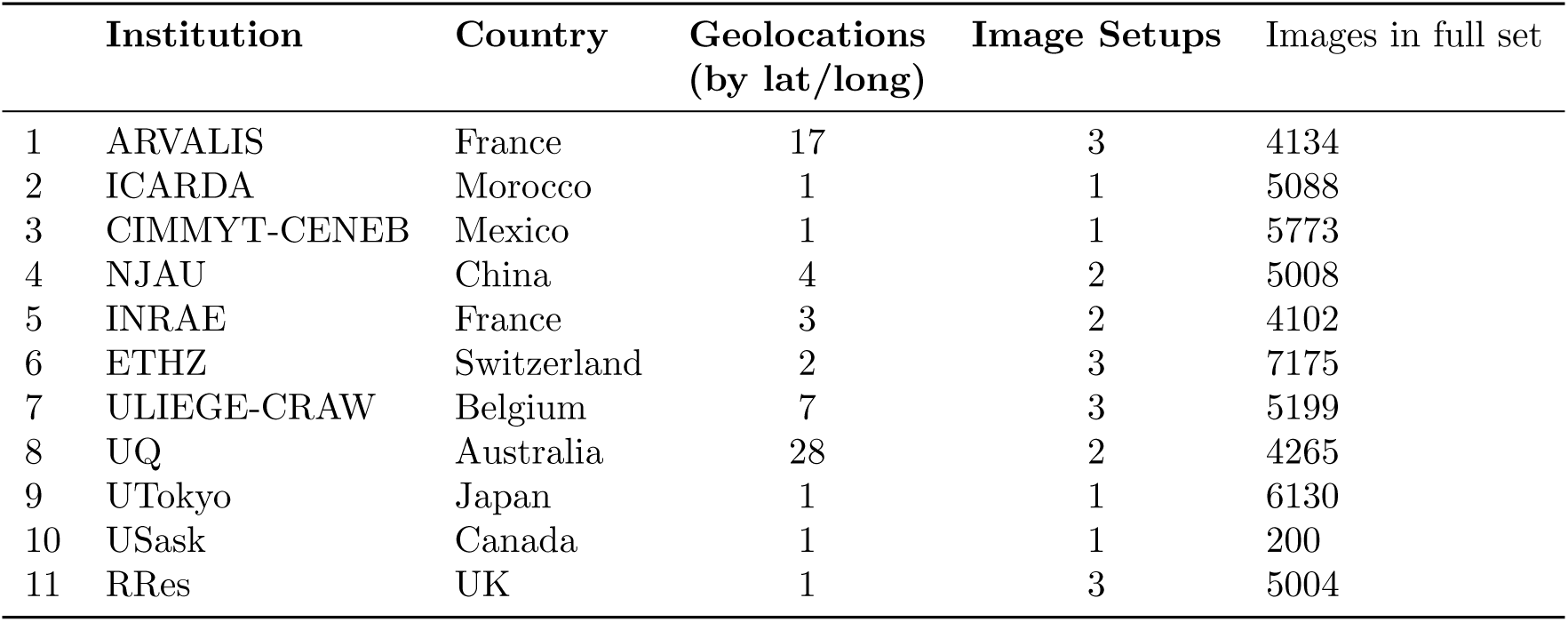
Summary of Institutions, the number of unique geolocations, the number of unique Image Setups, and the number of images per subset in the full dataset. A full description of the datasets (GWFSS v1.0 subsets.csv) is available in the ETH research collection referenced in the Data Availability section.

|  | Institution | Country | Geolocations<br>(by lat/long) | Image Setups | Images in full set |
| --- | --- | --- | --- | --- | --- |
| 1 | ARVALIS | France | 17 | 3 | 4134 |
| 2 | ICARDA | Morocco | 1 | 1 | 5088 |
| 3 | CIMMYT-CENEB | Mexico | 1 | 1 | 5773 |
| 4 | NJAU | China | 4 | 2 | 5008 |
| 5 | INRAE | France | 3 | 2 | 4102 |
| 6 | ETHZ | Switzerland | 2 | 3 | 7175 |
| 7 | ULIEGE-CRAW | Belgium | 7 | 3 | 5199 |
| 8 | UQ | Australia | 28 | 2 | 4265 |
| 9 | UTokyo | Japan | 1 | 1 | 6130 |
| 10 | USask | Canada | 1 | 1 | 200 |
| 11 | RRes | UK | 1 | 3 | 5004 |

**Table 2:** Imaging Setup Details. A full description of the imaging setups (GWFSS v1.0 imaging setups.csv) is available in the ETH research collection referenced in the Data Availability section.

| Imaging Setup | Vector | Camera Model | Viewing Angle | Focal Length (mm) | Sensor Resolution (pixel) | Field of View Horizontal (°) | Field of View Vertical (°) | Distance to Ground (m) | GSD (mm/px) |
| --- | --- | --- | --- | --- | --- | --- | --- | --- | --- |
| LITERAL1.0.0 | Handheld | Sony RX0 | 0 | 7.7 | 4800 x 3200 | 73 | 52 | 1.8 | 0.55 |
| LITERAL1.0.45 | Handheld | Sony RX0 | 45 | 7.7 | 4801 x 3200 | 73 | 52 | 1.2 | 0.32 |
| FIP1.0 | Gantry | Canon EOS 5D Mark II | 0 | 35 | 5616 x 3744 | 54.43 | 37.85 | 3 | 0.5495 |
| GO1.0M_0 | Cart | JAI GO-5000C-USB | 0 | 16 | 2560 x 2048 | 44.3 | 33.6 | 1 (to canopy) | 0.3125 |
| GO1.0M_30 | Cart | JAI GO-5000C-USB | 30 | 16 | 2560 x 2048 | 44.3 | 33.6 | 1 (to canopy) | 0.3608 |
| GO1.6M_0 | Cart | JAI GO-5000C-USB | 0 | 16 | 2560 x 2048 | 44.3 | 33.6 | 1.6 (to canopy) | 0.5 |
| Rres_GT3300_top | Gantry | Prosilica GT3300C | 0 | 50 | 3296 x 2472 | 38 | 26 | 2.8 | 0.12 |
| Rres_GT3300_south | Gantry | Prosilica GT3300C | 30 | 50 | 3296 x 2472 | 38 | 26 | 1.75 | 0.09 |
| Rres_GT3300_north | Gantry | Prosilica GT3300C | 30 | 50 | 3296 x 2472 | 38 | 26 | 1.75 | 0.09 |
| PhenoArm | Handheld | Sony RX0 | 0, 45 | 7.7 | 4800 x 3200 | 81.2 | 59.5 | 2 | 0.7143 |
| Phenotypette | Cart | Sony RX0 | 0, 45 | 7.7 | 4800 x 3200 | 81.2 | 59.5 | 2 | 0.7143 |
| Low-cost phenomobile | Cart | Canon EOS 600D | 0 | 55 | 1920 x 1080 | 23 | 15 | 2.4 | 0.046 |
| PHENOMOBILE | Ground Vehicle | Baumer HXG40 | 0 | 25 | 2040 x 2040 | 28 | 28 | 1.8 | 0.43 |
| Phenobuggy | Tractor Fobro | Baumer VCXG-124C | 0 | 25 | 4096 x 3000 | 32 | 24 | 1.8-2.4 | 0.24-0.33 |
| UFPS | Ground Vehicle | FLIR Chameleon3 USB3 | 0 | 16 | 2448 x 2048 | 26.4 | 19.8 | 2 | 0.45 |

1. **Arvalis** *ARVALIS 1-200* : The 200 subsets were acquired in 2022 and 2023 in a network of 18 sites representing the main agroclimatic zones and the most common practices in France. The trials cover different themes: evaluation of different wheat genotypes at diverse nitrogen fertilisation regimes, management methods, diseases, pests and water stress. Some trials comply with the specifications of organic farming or include wheat in combination with other species. Seed densities are a typical common practice in France. Images at most sites have been collected with the LITERAL [74], a handheld system with high-resolution cameras working without flash; The images at the location GROUX were acquired by PHENOMOBILE [9], an autonomous robot equipped with industrial cameras and flashlights.
2. **International Center for Agricultural Research in the Dry Areas (ICARDA)** *ICARDA MCH 2023* : Data collected at Merchouche Station (the main ICARDA experimental station near Rabat, Morocco), during a quite dry year. This set includes 960 entries from stage 2 of the durum wheat breeding program. Three acquisitions were taken with ICARDA’s PHENOBUGGY (equipped with RGB camera, multispectral and LiDAR) in March 2023. *ICARDA MCH 2024* : Data collected at Merchouche, also during a quite dry year. This set comprise 3 different trials (5 acquisitions each): 1) CWR panel as part of the Crop Trust project BOLD representing 60 elite durum wheat lines obtained from crosses with crop wild relatives; 2) CEREALMED including 288 entries of the Durum Global Panel of landraces, modern and old varieties mainly from Mediterranean countries; 3) a root rot trial of 24 elite lines.
3. **International Maize and Wheat Improvement Center (CIMMYT)** *CIMMYT-CENEB 1-7* : The CIMMYT dataset includes images collected for 319 spring wheat genotypes, consisting of elite, pre-breeding and exotic germplasm phenotyped during 2020 and 2021 field seasons in Campo Experimental Norman E. Borlaug (CENEB) in Ciudad Obregon, Sonora, Mexico. The genotypes were imaged from heading to maturity under irrigated, drought and chronic heat stress conditions in the field.
4. **Nanjing Agricultural University (NJAU)** The NJAU datasets were collected from field trials in different regions of China. Phenotyping data for *NJAU 1–NJAU 4* were collected using PhenoArm, a portable handheld imaging platform with two high-resolution cameras. NJAU 5 used Phenotypette, a pushcart platform integrating LiDAR, multispectral, and RGB cameras. The pushcart was manually operated at a controlled speed and equipped with RTK-GPS for automated data collection. *NJAU 1-2* : Experiments in Jurong and Xuzhou during 2020-2021, with 5 wheat cultivars under 3 nitrogen levels. *NJAU 3* : Trial in Xinxiang from 2022–2023 including 565 wheat cultivars, covering both introduced and domestic cultivars since 1950. Five cultivars were replicated 16 times, while the remaining 560 cultivars had no replication. A total of 640 plots were established, with fertilisation and irrigation managed according to local practices. *NJAU 4-5* : Trials in Yangling form 2021–2024 with the same cultivars as *NJAU 3*.
5. **National Research Institute for Agriculture, Food and Environment (INRAE)** The INRAE dataset was acquired in the frame of the FFAST project (French National Grant ANR-21-CE45-0037). The dataset includes images taken in field trials at three INRAE experimental sites UE APC at Auzeville (AUZ), UE DiaScope at Mauguio (MAU) and UE PHACC at Clermont-Ferrand (CLE) in the years 2021, 2022 and 2023. All pictures were taken using the Phenomobile V2 ground robot (https://hal.inrae.fr/hal-03646863), equipped with RGB cameras looking at nadir and at 45°. Images were taken in active illumination conditions (flashes). The trials consisted of 10 French elite cultivars grown under 4 treatments (depending on the site: irrigation, sowing date and seed density).
6. **Swiss Federal Institute of Technology Zurich; ETH Zurich (ETHZ)** *ETHZ 01* : Images from the ‘field phenotyping platform’ (FIP) at ETH Zurich in Eschikon [75]. The site covers typical climatic conditions of the Swiss Plateau. About 350 wheat varieties are monitored at least once per week to relate growth patterns to causal environmental factors. The set is available at [76]. *ETHZ 02* : Organic farming conditions at 981 m altitude, long snow cover and 2084 mm annual precipitation peaking in summer. Images show damage in spring caused by snow mould (Microdochium nivale). The high precipitation fostered lodging and diseases in the summer. There was high weed pressure.
7. **University of Lìege and Walloon Agricultural Research Center (ULIEGE-CRAW)** *ULIEGE-CRA-W 01-18* : Images were acquired in winter wheat trials in the Hesbaye area (Belgium) between 2018 and 2022. The 18 subsets detail the differences between the trials. Images cover mainly nitrogen fertilisation trials and nitrogen fertilisation × fungicide trials [10]. Images also cover N, P and K fertilisation trials (*ULIEGE-CRA-W 04*) and drought experiments (*ULIEGE-CRA-W 12, 18*). The set also contains sample images from dense time series of the same plots recorded in 15-minute intervals (*ULIEGE-CRA-W 11*). One series contains green reference spheres used as control points in thermal images acquired in addition to RGB images (*ULIEGE-CRA-W 17*).
8. **The University of Queensland (UQ)** UQ 1-29: The 29 subsets detail the differences between trials in which the images were acquired. Images are collected in the 2020 and 2021 National Variety Trials. Differences include variations in geolocation, genotype, and growth stage across Australia. The photographs were taken in 2020 and 2021, using smartphone cameras from a top-down perspective at about 0.5 to 1.5m above the canopy.
9. **University of Saskatchewan (USask)** The USask dataset was collected from wheat phenotyping field trials at the Kernen Crop Research Farm in Saskatchewan, Canada in 2019. The images comprise a single field trial with 32 diverse wheat cultivars at the heading stage. Images were collected with the University of Saskatchewan Field Phenotyping System (UFPS), a custom-built, self-propelled ground vehicle equipped with a range of imaging instrumentation, RTK-GPS, and on-board data processing.
10. **University of Tokyo (UTokyo)** The UTokyo dataset was collected from wheat phenotyping field trials at the Institute for Sustainable Agro-ecosystem Services (ISAS) in Tokyo, Japan, in the 2014-2015 season. A Field Server system [77] collected images of five genotypes through the whole growth stage. The camera module of the system is based on a digital single-lens reflex (DSLR) camera, the Canon EOS Kiss X5 camera, with an EF-S18-55 mm lens (Canon Inc., Tokyo) that provides high-quality and high-resolution (18 megapixels) image data. A preprogrammed microcontroller board controls the power and shutter of the camera automatically.
11. **Rothamsted Research (RRes)** The Rothamsted dataset includes images collected for 391 wheat genotypes, captured throughout the growth cycle, from tillering to maturity, using the Field Scanalyzer [11]. The NIT subsets relate to the evaluation of four commercial variety growing supply with six levels of nitrogen input over two years (2019 and 2021). Images of the NIT subsets were captured from three angles: 30° north, 0° top, and 30° south, providing comprehensive spatial coverage. The PxCS and the PxG subsets provide images from two mapping populations that were planted in 2019 and 2021, respectively. The populations displayed a large range of variation in terms of phenology and height.

### 2.2 Image Acquisition

The imaging setups consist of a vector and a camera ranged from hand-held over manual push-cart to fully automated rovers, gantry systems or a cable-suspended system mounted on poles (Table 2). Images were acquired with RGB cameras of at least 1920 x 1080 pixel sensor resolution, oriented from nadir (0°) to 45° viewing angle. All carriers positioned the camera between 3 and 1 m above the ground, leading to ground sampling distances between 0.09 and 0.71 mm.

### 2.3 Data Selection for the Annotation Pool

For data selection, we entrusted expert judgment. To assemble a diverse set of images for annotation, each participating institution was asked to provide approximately 5,000 images encompassing different phenological stages for the data set and a diverse subset of 200 for the annotation pool. The selection of images prioritised diversity across key factors, including variations in phenological stages, geographic locations, cultivars, and imaging conditions. Additionally, institutions were encouraged to include treatments, such as nitrogen levels, irrigation, or variation in other inputs, aiming to encompass a broad spectrum of scenarios in the dataset. From the annotation pool, a total of 1,096 images were selected through a stratified approach, ensuring proportional representation across contributing institutions. Specifically, the selected image set comprises 110 images from each of the seven institutions (*i.e.*, INRAE, ETHZ, USASK, Arvalis, RRES, NJAU, and UQ), 109 images from two institutions (*i.e.*, ULiege and Utokyo), and 108 images from CIMMYT.

The selection process was driven by a joint consideration of feature geometry and institutional balance. Specifically, image features were extracted using a ResNet model [78] pre-trained on ImageNet. A k-means clustering algorithm was then applied to group the images based on their feature similarity. To ensure representative sampling, images closest to the cluster centres were selected, while also maintaining a balanced distribution across institutions. This process was designed to maximise the uniqueness of the selected images based on their embedding distributions. As a final data preprocessing step, all images were standardised to ensure consistent resolution and comparable ground sampling distances. This was accomplished by applying a centre crop to achieve a resolution of 512 × 512 pixels for most datasets. An exception was made for data contributed by UTokyo, where a resolution of 1024 × 1024 pixels was used to accommodate the more detailed ground sampling distance. Figure 2 illustrates the images sampled from our proposed dataset, showcasing the diversity of the proposed dataset.

**Figure 2:**
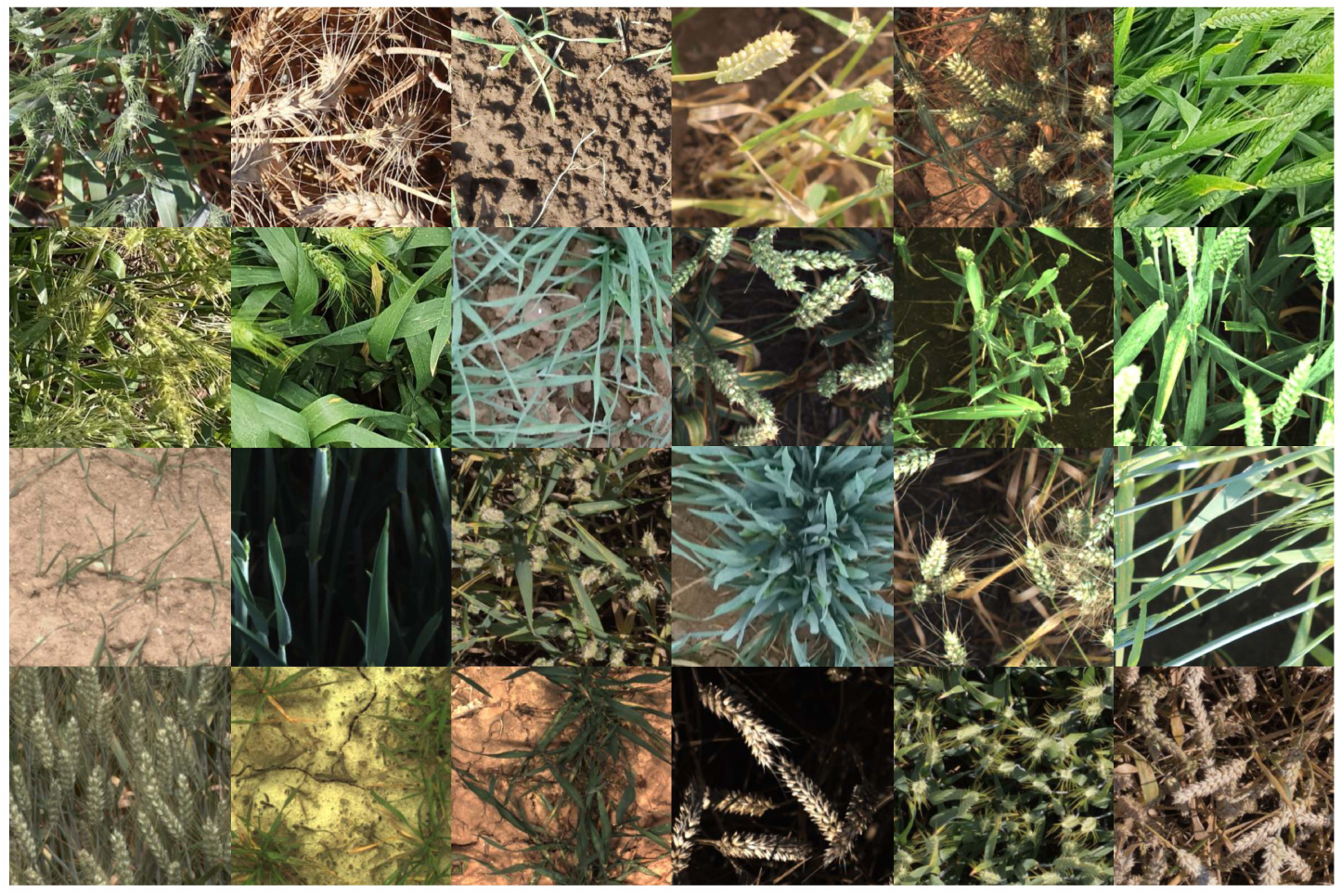
Representative samples from our proposed GWFSS dataset.

### 2.4 Labeling

#### 2.4.1 Targeted wheat entities

The annotation process was carried out centrally by expert annotators using the Darwin annotation tool provided by V7 Darwin ^1^. During annotation, temporary adjustment of brightness and contrast was done to enhance the distinctions among features. The annotation process and quality control were handled by HIPHEN. In case of annotation mistakes, images were sent back to the annotators with respective instructions. In these cases, segmentation masks were modified using the brush and eraser tools. The global wheat experts team (Table 7, labelling) reviewed and resolved the unclear cases as needed. We refer readers to the appendix for the detailed GWFSS labelling guide. Initially, a small set of image tags was assigned to enhance understanding of image content and quality (see Table 3). Subsequently, pixel-level annotations were performed for the following classes: head, leaf, stem and background. As a reference for tissue types, we largely used the BRENDA Tissue Ontology (BTO^2^) retrieved in the EMBL-EBI Ontology Lookup Service ^3^. The targeted entities were i) “heads” defined as spike (BTO 0001278) excluding awns (BTO 0005641), ii) “leaves” defined as the leaf lamina (BTO 0000719) including ligule, and iii) “stems” (BTO 0001300) including the surrounding leaf sheath (BTO 0005094). The peduncle was not labeled separately but included in “stem” (Fig. 3). The peduncle (PO 0009053) is the shoot axis that extends from the last foliage leaf on a stem (*i.e.* the flag leaf) until the next distal node (*i.e.* the basal end of spike).

**Figure 3:**
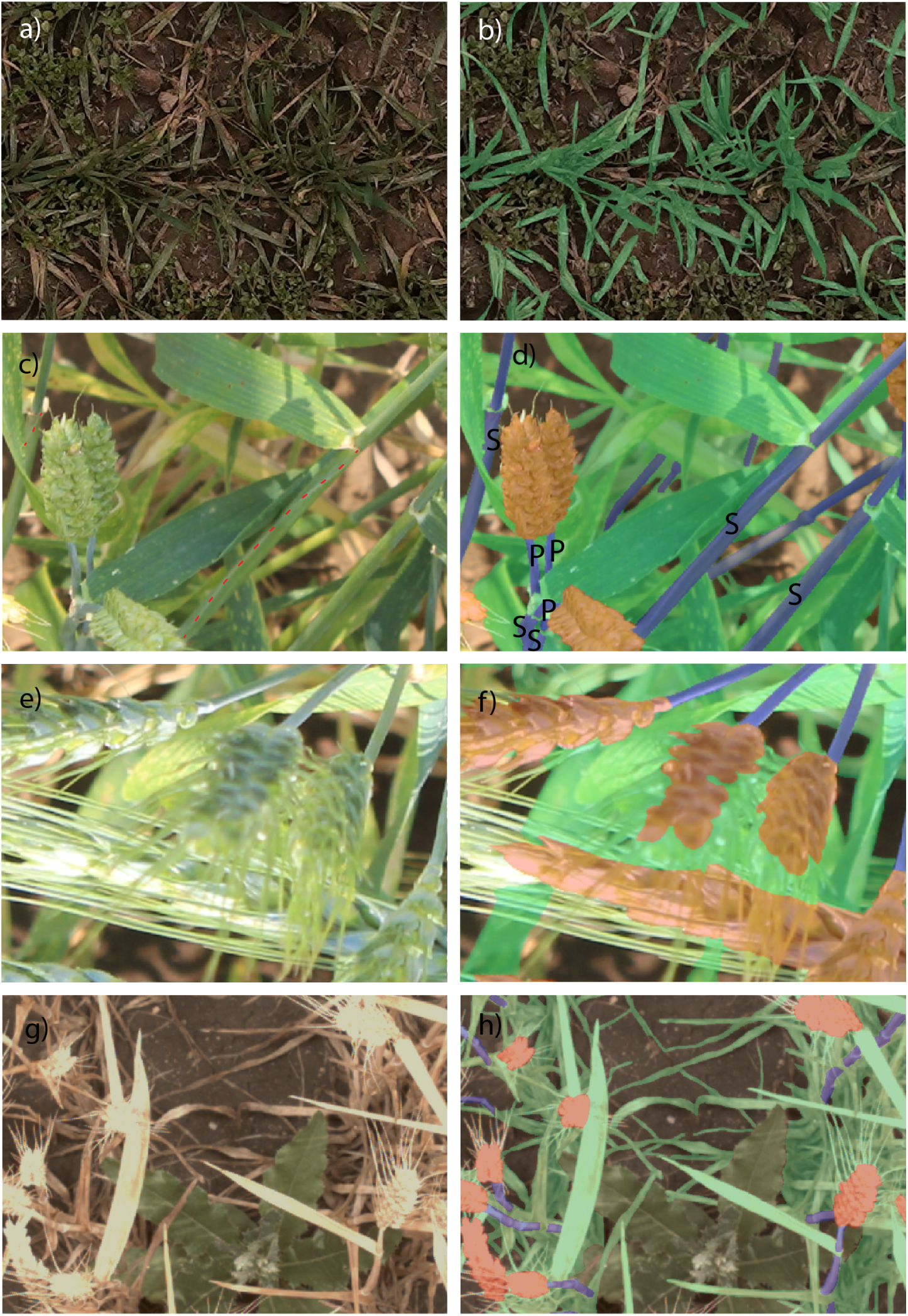
Examples for the labelling process with original images (a, c, e, g) and labelled images (b, d, g, h) showing the segmentation masks for leaves (green), stems (purple) and spikes (orange). Red dotted lines in a) show the edge of the leaf sheath wrapped around the stem. A leaf sheath (S) as part of the stem can be either recognized by this edge or its connection to a leaf blade. A peduncle (P) is a part of the stem located between a visible spike and the collar of the flag leaf. For our analysis awns were treated as invisible by drawing the segmentation masks above them (f, h). Weeds and crop residues were considered background (b, h)

**Table 3:** Imaging Tagging Overview.

| Tag Name | Description | Possible Value |
| --- | --- | --- |
| Institute | The institute that contributes this image. | INRAE, ETHZ, USASK, Arvalis, RRES, NJAU, UQ, ULiege, Utokyo, CIMMYT. |
| Name | The unique image name. | Not Applicable. |
| Size | The width and height of the original image. | (4800×3200), (2048×2048), (4096×3000), (3456×5184), (5184×3456), (5634×3753), (2040×2044), (4080×2704), (4080×3200), (3296×2472), (2560×2048), (1024×1024), (5184×3456). |
| Anthers | The existence of anthers. | True, False. |
| Bending | The existence of ear bending. | True, False. |
| Lighting | The existence of shadow. | True, False. |
| Stage | Phenological Stage. | Emergence, Vegetative, Stem Elongation, Ear Emergence, Early Filling, Early Senescence, Late Senescence, Maturity. |

Consequently, the wheat stem labelling included leaf sheaths, peduncles and bare stems (e.g. towards the end of the growing season). This decision considers that it is difficult to separate the different classes in complex images. The clear identification of the peduncle requires a visible ear (indicated by “P” in Fig. 3 d); the clear identification of a leaf sheath requires that either its edge or the attached leaf blade is visible (indicated by “S” in Fig. 3 d). In many cases, it was not possible to decide if the structure was sheath, bare stem or peduncle. The wheat head label specifically encompassed only spikelets, excluding awns. Awns are distal bristle-like extensions of the lemma surrounding the florets of wheat [79]. Thus, while spikelets including glumes and florets were labelled as part of the wheat head, awns were treated as “invisible” features (Fig. 3 f, g). There are practical challenges in annotating individual awns in all imaging scenarios, particularly when awns appear blurry or lack distinct contours. Thus, when awns overlapped with the targeted organ, they were treated as if they were absent and the polygon was drawn across the organ in the background (Fig. 3 g, f). Detached senescent plant material, such as debris or residue resulting from no-till practices, was excluded from annotation and classified as background. All targeted entities were labelled as long as they were still attached to the wheat plant regardless of their colour, *i.e.* chlorotic or necrotic tissue was also labelled.

The annotation process was exclusively focused on wheat, with weeds deliberately left unannotated and classified as background. All other non-wheat objects were similarly disregarded and annotated as background (Fig. 3 b, h).

### 2.5 Baseline Segmentation Models Development

**DeeplabV3Plus** [80] is a classic semantic segmentation framework based on convolutional neural networks (CNN) that employs an Encoder-Decoder architecture. The model builds upon the strengths of Deeplabv3, which leverages Atrous Convolution to explicitly control the resolution of feature maps and adjust the reception field. In Deeplabv3Plus, encoder features are first up-sampled bilinearly by a factor of 4 and then concatenated with the corresponding low-level features from the backbone network. 1×1 convolutions are applied to low-level features, reducing the number of channels to reweight rich contextual encoder features and simplify training. After the concatenation, the model refines these combined features using a series of 3 × 3 convolutions, ensuring the integration of detailed spatial information and high-level semantic context. The decoder finalises the segmentation mask with a simple bilinear upsampling operation by a factor of 4, delivering high-resolution predictions. This seamless combination of multiscale context aggregation through the encoder and spatial detail recovery in the decoder positions Deeplabv3Plus as a robust and flexible solution for semantic segmentation tasks.

**Segformer** [63] processes an input image of size *H×W ×*3 by first dividing it into 4×4 overlapping patches. These patches are fed into a hierarchical Transformer encoder to extract multilevel features, leveraging an Overlapped Patch Merging strategy to ensure spatial continuity and capture richer local context. The encoder utilises efficient self-attention mechanisms by applying dimension reduction, which reduces the time complexity of the self-attention mechanism. This significantly optimises computational efficiency with minimal impact on segmentation performance.

The extracted features are then processed by a lightweight multi-layer perceptron decoder. Unlike traditional hand-crafted designs such as Deeplabv3Plus, this approach simplifies the decoding process while improving the effective receptive field, enabling precise and efficient segmentation. To cater to diverse performance and resource requirements, Segformer introduces a family of Mix Transformer encoders (MiT-B0 to MiT-B5), which share the same architecture but vary in size, offering flexibility in balancing computational cost and segmentation accuracy. By integrating innovative design choices with practical adaptability, Segformer delivers a robust and efficient solution for semantic segmentation tasks.

#### 2.5.1 Impact of Distribution Shift on Segmentation Performance

We conduct experiments under two different data-splitting settings. (1) **Random Split:** In this setting, we randomly split the data into a training set (70%), a validation set (10%) and a test set (20%). (2) **Region Split:** In this setting, we utilised data from Arvalis, CIMMYT, ETHZ, INRAE, NJAU, RRES, and ULiege CRA-W as the training set, data from UTokyo as the validation set, and data from UQ as the test set. The UQ test set is rather challenging due to the massive diversity in genotypes and Australian growing environments and imaging conditions. For both of the settings, the validation mIOU was used to select the best checkpoint, which was then used for testing.

#### 2.5.2 Impact of Training Data Scale and Model Size on Segmentation Performance

We conducted two sets of experiments to investigate how the size of the training dataset and the number of model parameters influence the performance of the segmentation. To investigate the relationship between the size of the training dataset and the performance of the model, we trained SegFormer-B0 using progressively larger subsets of the full training dataset. Specifically, we sampled subsets containing 1%, 5%, 10%, 20%, 30%*, · · ·,* 100%, of images from the full training set to train the model. To assess the impact of model size on segmentation performance, we trained SegFormer-B0 to B5 using the full training dataset. As the model progresses from B0 to B5, both the number of parameters and computational cost increase, allowing us to analyse how model complexity affects segmentation accuracy.

### 2.6 Evaluation metric

**Mean Intersection over Union (mIoU)** measures the level of overlap between the predicted mask and the ground truth mask. Specifically, we have:

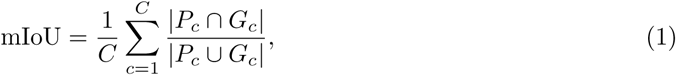

where *P_c_*and *G_c_* denote the predicted mask and ground truth mask of the *c*-th class.

**Mean Pixel Accuracy** focuses on the pixel-wise accuracy for each class, which can be defined as:

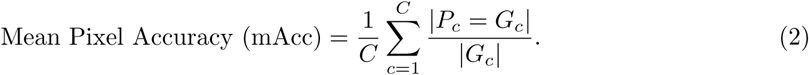

## 3 Results

### 3.1 UMAP Visualisation of Image Diversity

The diversity of the training dataset is critical for the generalisation capacity of segmentation models. The diversity of a crop dataset can be affected by multiple factors, such as differences in phenological stages, lighting conditions, background, growing environment conditions, and genotype. In this work, the distribution of GWFSS images was analysed using the UMAP technique on image features extracted by an ImageNet-pretrained ResNet-50 model. The image embeddings of the top two UMAP components visualise the distribution of all labelled GWFSS data. The visualisation of images in the latent space was either colored by the phenological stage (Fig. 4 a) or institution (Fig. 4 b). For the phenological stages, the first UMAP dimension shows a clear clustering in the sequence of stage progression. With the first UMAP feature increased from 0 to 13 (Fig. 4 a, dimension 1), the growth stage generally progresses from emergence and vegetation towards senescence and maturity. Most of the images taken from emergence to stem elongation clustered at a value below 7, while the other extreme images around late senescence and maturity clustered above 7. Images containing ears, i.e., starting from ear emergence, showed values above 4 in dimension 1. The clustering in the second dimension tended to be driven by the institution providing the images. Images from USASK, ULiege, and RRES generally have the second UMAP feature valued below 8, while most of the UQ and NJAU features are above 8. By viewing the images of these institutions, this phenomenon could be attributed to the difference in the lighting conditions. The datasets of UTokyo and NJAU cover a wider range of dimension 2, while the dataset of Arvalis is the only one spanning almost the entire latent space.

**Figure 4:**
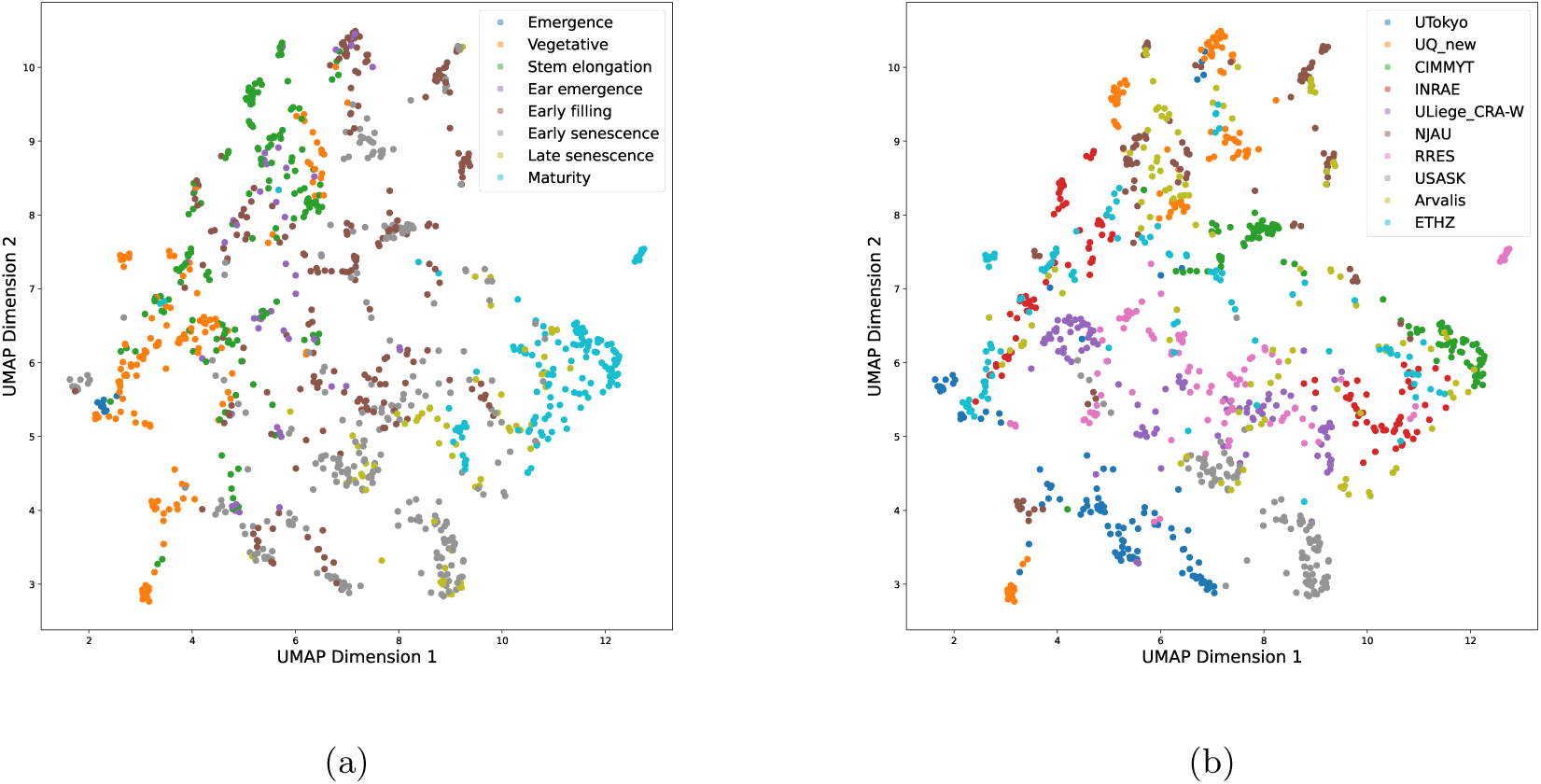
The UMAP visualisation of GWFSS labelled images coloured by (a) phenological stage and (b) institution.

### 3.2 Balance of Developmental Stages and Labelled Classes

Images were tagged with the approximate developmental stage estimated from the image (i.e. stages were not recorded as ground truth in the field). Although the aim was to balance all the phenological stages, this was not possible for all datasets. An analysis of the tags across the whole dataset revealed an uneven distribution across different phenological phases (Fig. 5, a). Notably, ‘early filling’ and ‘early senescence’ were the most frequently observed (224 and 282 images, respectively), while ‘emergence’ was the least represented stage (8 images).

**Figure 5:**
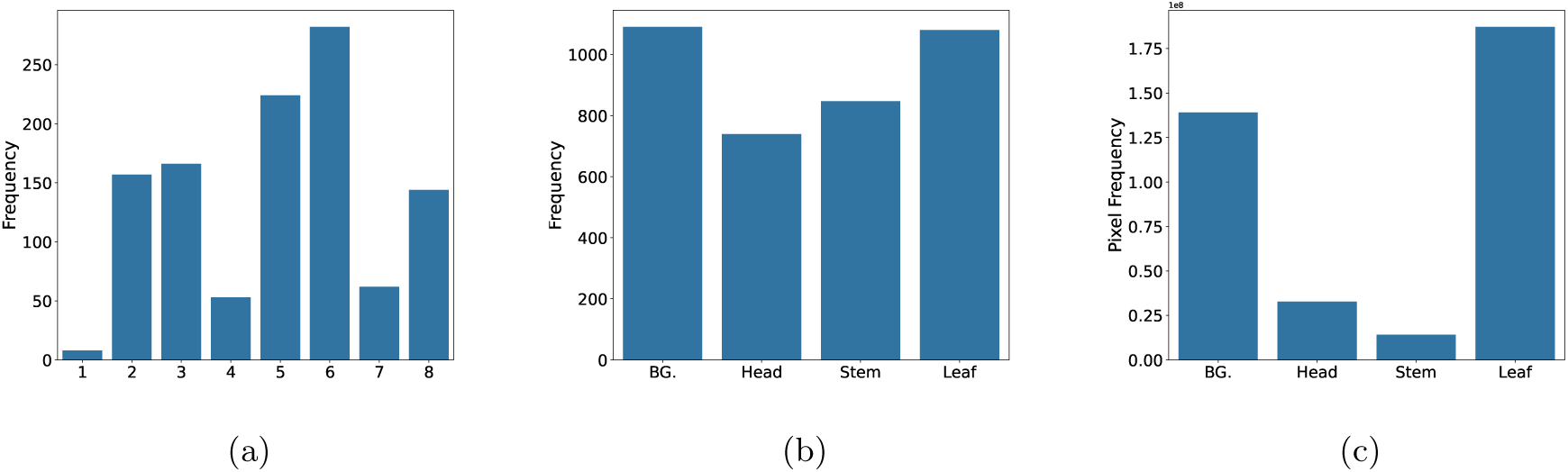
Statistics of the GWFSS dataset: (a) The distribution of growth stages for each image, where 1 *−* 8 on the x-axis indicates ‘Emergence’, ‘Vegetative’, ‘Stem elongation’, ‘Ear emergence’, ‘Early filling’, ‘Early senescence’, ‘Late senescence’, and ‘Maturity’, respectively. (b) The class occurrence at the image level. (c) The class occurrence at the pixel level.

We also evaluated the balance of the labelled classes. At the image level, background (BG) and leaves were the most prevalent, appearing in 1,090 and 1,080 images, respectively (Fig. 5, b). In contrast, the stems and heads were present in 847 and 739 images, respectively, since they predominantly emerge in the later phenological stages. At the pixel level, a more pronounced class imbalance was evident, with the leaves occupying the largest proportion of pixels, followed by the background, while the stems and heads account for significantly fewer pixels (Fig. 5, c). Thus, although heads and stems appear frequently at the image level, they still constitute only a small fraction of the total pixel distribution, compared to leaves and background.

### 3.3 Baseline Segmentation Models

Concerning the sampling strategies to split images into training, validation and test sets, there was a noticeable performance gap between the random split vs region split strategy (Table 4). We attribute this discrepancy to the distribution shift between the training and test data in the region split setting. Among the evaluated models, Segformer consistently outperformed DeepLabV3plus, achieving a 2.8% higher mIoU in the random split setting and an 8.5% improvement in the region split setting. Notably, in the Region split strategy, Segformer performed substantially better for Head and Stem classification than did DeepLabV3plus with little difference in estimation of Background and Leaf. These results highlight the superior capability of Segformer in addressing the wheat organ segmentation task, especially under distribution shifts.

**Table 4:** Comparison of Mean Intersection over Union (mIoU) and mean accuracy (mAcc) metrics for Deeplabv3plus (R101) and Segformer (B1) across Random- and Region-based data splittings.

|  | Segformer (B1) |  |  |  | DeeplabV3+ (R101) |  |  |  |
| --- | --- | --- | --- | --- | --- | --- | --- | --- |
|  | Random |  | Region |  | Random |  | Region |  |
|  | IoU | Acc | IoU | Acc | IoU | Acc | IoU | Acc |
| **Background** | 84.51 | 92.75 | 75.46 | 88.71 | 81.59 | 89.88 | 74.32 | 82.11 |
| **Head** | 82.85 | 90.14 | 66.11 | 86.84 | 81.46 | 89.05 | 46.25 | 48.27 |
| **Stem** | 44.92 | 53.89 | 19.23 | 20.95 | 39.40 | 46.73 | 6.64 | 7.06 |
| **Leaf** | 82.35 | 90.44 | 81.75 | 89.36 | 80.93 | 90.81 | 81.4 | 95.14 |
| **Average** | 73.66 | 81.81 | 60.64 | 71.47 | 70.85 | 79.12 | 52.15 | 58.15 |

Concerning the size of the training data, the overall model performance sharply increased between 1% and 60% (i.e. 460 images) of the training data and plateaued thereafter (Fig. 6, a). Above these 460 images, there were only marginal improvements as the dataset size approached 100%. The heads and leaves were well segmented with only 10% of the data (*IoU >* 75%) and the model performance only progressed slowly when more data were used. The segmentation of stems improved with increases to 60% of the training data but the performance plateaued at a low IoU of 40%.

**Figure 6:**
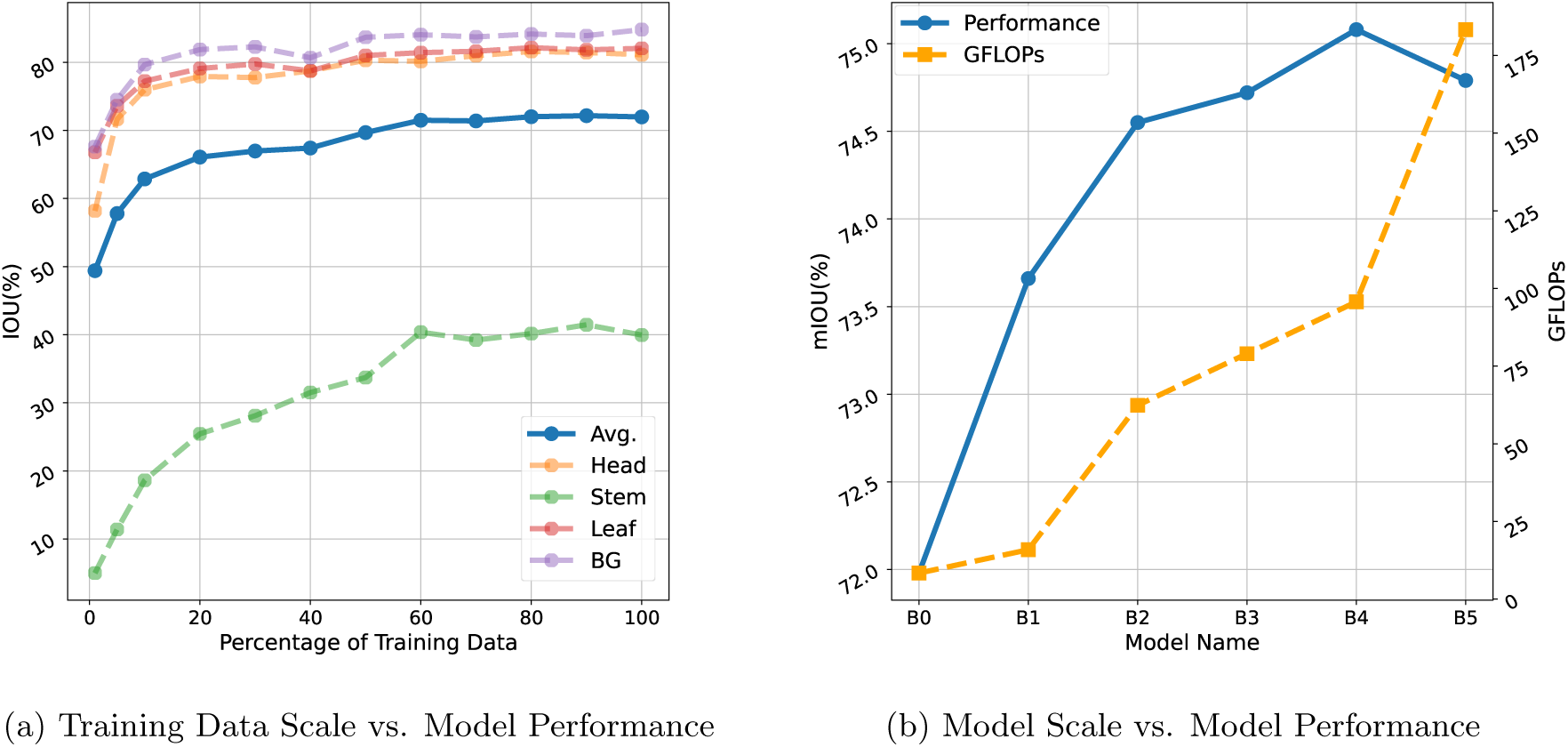
Effect of (a) training data size and (b) segmentation model size on Segformer model performance. The effect of training data size on the intersection over union (IoU) was evaluated for all object classes including the background (BG). The segmentation model size performance was evaluated as mean IoU (mIoU) and Giga Floating Point Operations per Second (GFLOPS). The higher GFLOPS indicates heavier computational complexity. Here B0 indicates the smallest Segformer model while B5 indicates the largest Segformer model.

To examine the impact of model size, we trained SegFormer models of varying capacities (B0 to B5) on the full training dataset and analysed their performance (Fig. 6, b). The Giga Floating Point Operations per Second (GFLOPS) served as an indicator of computational complexity, where higher values denote greater computational demands. The results indicate a steady improvement in performance from B0 to B4. However, when training SegFormer-B5 on the full dataset, performance degradation was observed. We attribute this decline to the insufficiency of training data to adequately support the significantly larger parameter space of SegFormer-B5, which nearly doubles that of B4.

### 3.4 Visual Inspection of Segmented Images

Key requirements of the model were to detect organs independently of their colour and distinguish them from weeds or plant residues on the ground. We conducted a systematic review of all five images that contain weeds in the test set (Random Split, as described in Section 2.5.1). The weeds were of different types with different leaf shapes and were all classified as background (Annex, Fig. S2). A sample is displayed in Fig. 7, a, k and l. Late in the season, weeds frequently germinate between rows and obscure the senescence signal. Similarly, the separation of plant residues from wheat plants (Fig.7, b) is an important advantage. When awns are present, the segmentation of the spike without the awns (Fig. 7, a, d, f, h, i, j, k) is a useful feature that can assist in the approximation of spike volumes. To evaluate the segmentation of senescing canopies, we selected images from early senescence to maturity within the random split test set (as detailed in Section 2.5.1). This set representing later stages comprises 73 images. In general, the model maintains strong performance (Table S2). However, performance declined relative to the full test for the background (IoU of 73.5 vs. 84.5 for late stage vs. full set) and leaves (IoU of 73.7 vs. 82.3 for late stage vs. full set). This is likely due to the increased visual similarity between senescing leaves and background elements (e.g., soil or dried residue), as well as reduced structural distinctiveness in ageing foliage, making accurate segmentation more challenging.

**Figure 7:**
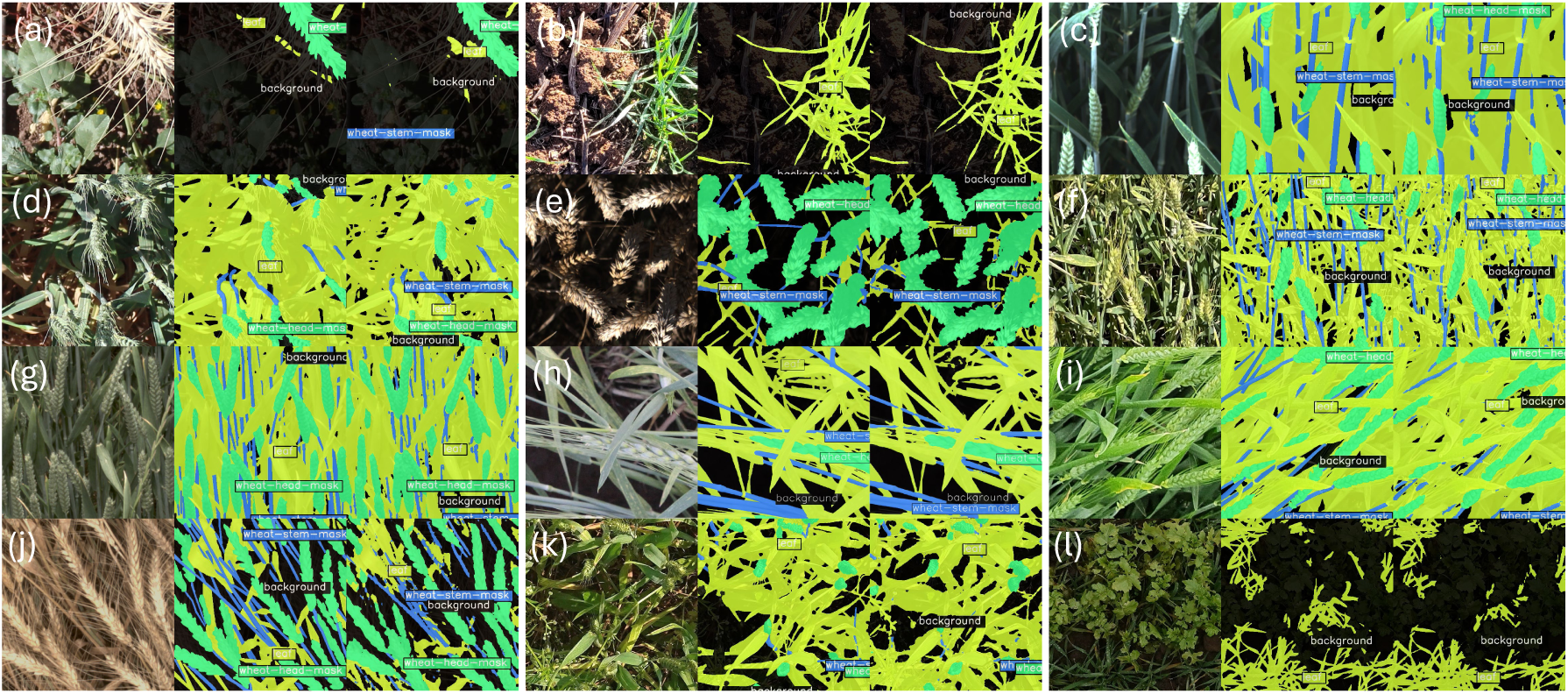
Visualisation of original image (left strips), ground truth (centre strips) and prediction results (right strips) from Segformer-b1

## 4 Discussion

### 4.1 The Challenges of Organ Labelling in Complex Canopies

We collected 52,078 images from 67 different field sites worldwide with a ground sampling distance (GSD) between 0.09 and 0.71 mm per pixel. The UMAP visualization confirms the need for such diversity, as it shows clustering by institution and developmental stage.

The collaborative effort to sample this diverse set of images and to design the labelling strategy was essential to the success of the work. All participating institutions operate imaging setups collecting images of wheat canopies in the sub-millimeter range and are at the forefront of enhancing the in-depth analysis of complex canopies. A first step was the decision of which canopy features could be targeted, given the available spatial resolution. In wheat, ground sampling distances below 10 mm permit good estimates of canopy cover and leaf area index or crop density, while GSDs below one mm are needed to detect individual leaves of emerging seedlings [81]. The given sub-millimeter resolution was sufficient to label the targeted organs. Reliable labelling of awns would likely require GSDs below 0.2 mm and substantially more labour for manual labelling. New sensors with higher resolution will allow for awn segmentation, even when operating with the same carrier system.

Given the experience in GWFSS, the term “plant organ” needs to be understood in the context of imaging constraints. With this regard, we would like to add a “sidenote” to protocols for the minimal requirements to describe plant phenotyping experiments. The MIAPPE 1.1. [82] release states that *“Observed variables, traits, methods and scales are each identified by name, and may have a reference to the corresponding ontology concept (ideally from the Crop Ontology)”*. However, in the Crop Ontology database^4^, entities upon which traits are measured are not always indexed with their own identifiers. For example, while stem colour (CO 321:0000973) is defined as “colouration of the stems,” the entity “stem” itself is not defined. GWFSS aims to extract organs as entities on which traits will be measured. For this reason, we prefer the BRENDA Tissue Ontology (BTO^5^), which is a vocabulary for the source tissues. As such, it focuses on a detailed description of the entity rather than the trait. The segmentation of the entities is the first step towards deriving new phenotypes. Moreover, our digitally extracted “organs” do not quite comply with classic ontology terms: heads exclude their spikes, and stems include leaf sheaths. One challenge is therefore how to define a trait which is based on more than one organ part (*e.g.* stem = true stem plus leaf sheath), given that trait ontologies are typically structured hierarchically. Along these lines, Celestina et al. [83] have identified the need to reconsider the classical growth scales, such as Zadoks [84] or the derived unified BBCH scale [85] to fit the needs of image-based phenotyping. They draw up a list of the development stages that need to be assessed by destructive sampling and the stages that can be assessed in a non-destructive manner. We believe that their phases of the Population of Culms Development Scale (PCDS) may be enhanced by image-derived phases. At least heading and physiological maturity can be digitally measured based on models derived from GWHD and GWFSS as we will discuss below.

### 4.2 GWFSS Compared to Other Datasets for Semantic Segmentation of Wheat Organs

The availability of extensive open datasets has been crucial to remarkable progress in applications of modern computer vision in agriculture. Various datasets have been released to facilitate computer vision applications, such as for fruit detection [86], weed management [87], green coverage estimation [88], and plant disease identification [89]. For semantic segmentation, pixel-level annotation remains a cornerstone of segmentation tasks, but it is notoriously labour-intensive and expensive. Intensive collaboration among public and private institutions is needed to generate sufficiently large, diverse, and consistently labelled data.

The novelty of the GWFSS dataset is its diversity, its coverage of all stages, and labelling of all organs of wheat. The GWFSS dataset contains fewer annotated images than GWHD [16], which labelled 6,510 images from 16 institutions. However, GWFSS dataset samples the whole growing season while GWHD focuses on head detection during flowering, grain filling, and ripening. Moreover, for the GWFSS dataset, we decided to supply a large set of 52,078 images to enable users to pose solutions to other questions by applying their labelling. A similar dataset assembled by the global-rice dataset consortium is underway for rice [73]

For semantic segmentation, there are several smaller datasets available which are not included in GWFSS, mainly because they focused only on specific organs of wheat. Although most studies trained segmentation models, we will summarise only the characteristics of the annotated datasets as the most valuable part of the studies (see Table 5). With regard to wheat heads, several smaller datasets were collected in addition to the GWHD dataset [36, 65, 66]. Few researchers have collected datasets to train leaf segmentation during the early [72] or late phases of development [73]. The latest trend is to target features within organs within complex canopies in the field, such as spikelet segmentation [90], Fusarium head blight [71], or leaf diseases [91]. For such annotations of wheat head damage, a concise and coordinated labelling of diverse datasets may be highly valuable.

**Table 5:**
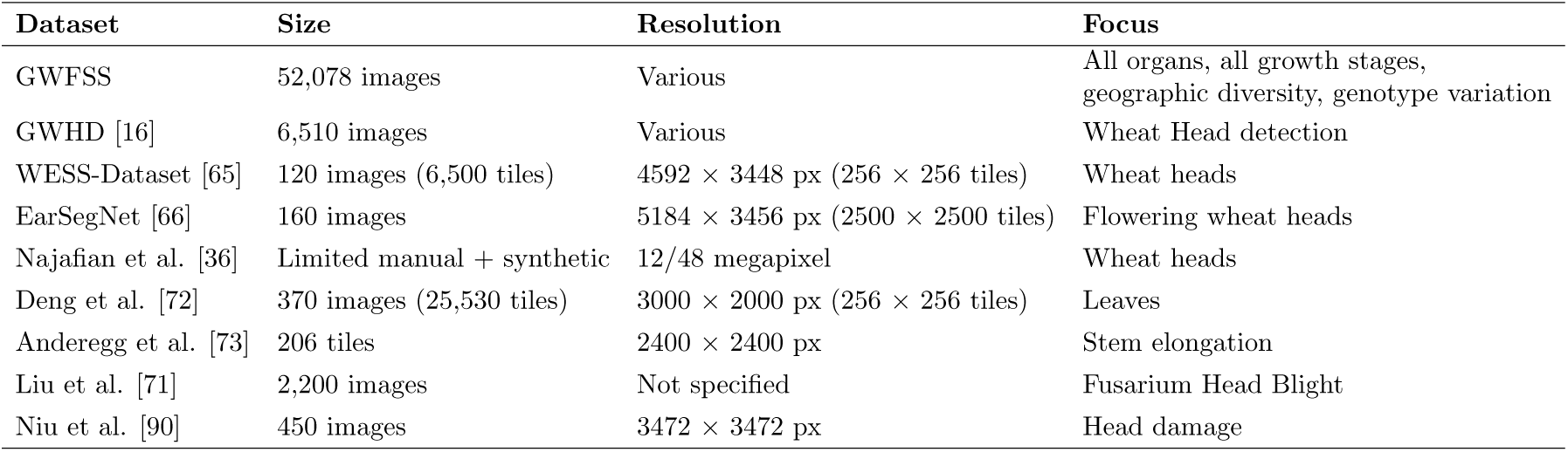
Summary of wheat segmentation datasets.

### 4.3 Organ Segmentation - a path to integrative traits

Field phenotyping is often considered in the context of spatial and temporal scales. We believe that the ability to track organ development throughout the growing season will set a new standard for phenotyping. It will aid breeders, variety testers, or researchers in evaluating genotypes for improved canopy architecture, source-sink balance and resistance to environmental and disease stressors.

The ability to follow the different wheat organs through the season enables the testing of more complex phenotypes. For example, our model would permit quantification of the changes in tissue reflectance as they change from green to chlorotic to necrotic. In wheat, this was done, for example, using shallow learners based on colour spaces using a support vector machine classifier [19] or a multiclass random forest classifier [73]. Canopy segmentation followed by reflectance indices or the mentioned classifiers opens new possibilities for field phenotyping. For example, changes in leaf colour can be used to quantify canopy damage in winter due to frost or diseases. Later, it may be used to quantify nitrogen status.

Stem (or more precisely peduncle) senescence is a measure of physiological maturity of wheat [92]. It is time-consuming to assess and, therefore, rarely reported. Using a small semantic segmentation training dataset, Anderegg et al. [43] tracked the senescence process of leaves, stems, and heads through grain filling and showed that stems were the last organ to be senescent. In this work, the genotypes differed in their dynamics and timing of leaf, head, and stem senescence, highlighting the relevance of tracking the organs separately. The GWFSS training dataset is a major step forward in measuring physiological senescence and separating different canopy senescence processes.

We acknowledge that proximal sensing is currently not the primary choice in cases where thousands of plots are to be evaluated and breeding programs are more likely to require high-throughput remote sensing by means of unmanned aerial vehicles (UAV). Remote sensing by UAV equipped with lower pixel resolution multi-spectral sensors allows the estimation of traits like LAI and canopy senescence. However, these sensors are not ideal to study the different organs in the canopy. Plant organs may have substantially different proportions in the canopy, influencing its reflectance spectrum. Variation with time occurs due to environmental effects on crops (leaf rolling, frost, heat, drought, nutrition, and pest effects), stem elongation, spike appearance, and bending or lodging of spikes during grain filling. The transferability of reflectance spectra from one season or trial to a different one may be influenced by environmental conditions or diseases. Although proximal sensing has lower throughput, it can complement aerial measurements with information from the organ scale. This might enable upscaling from proximal to remote sensing. Alternatively, high-resolution RGB imaging is becoming readily available on UAV with cameras with a resolution ranging from 60 to 120 Mpx, though UAV are slow and difficult to localise when flown close to plots to take still shots. A solution is to fly UAVs closer to the plots and use video at high-shutter speeds to avoid blurring the image or disrupting the canopy with propeller downwash [93] were able to achieve a GSD of 0.13m flying a 20 Mpx camera using video at ca. 5m above the canopy at a speed of 2-3 m/s. Finally, there is an increasing number of phenotyping robots available, as well as higher resolution RGB UAV cameras that allow organ phenotyping of larger numbers of plots.

We recognise that RGB datasets are only a piece of the larger phenotyping toolbox: Other technologies could and should be used in combination with data fusion methods. For example, LiDAR has been used to estimate biomass and crop growth rate [94]. Future RGB datasets collected at the same time as LiDAR point clouds can be used to study biomass accumulation at deeper hierarchical levels, such as biomass partitioning in the different organs. Furthermore, RGB combined with thermal imagery can be used to assess abiotic stresses [95]. Breeders could focus on targeted breeding for specific organs and select new genotypes with increased water use efficiency, to exemplify a few cases where RGB organ segmentation can be made the most of.

### 4.4 Baseline Model to Guide the Size of the Training Data

The main contribution of this work relates to the creation of a large open-access database of images capturing diverse field-grown wheat plots. However, without a basic model it was difficult to judge how large the training data should be. According to the trained baseline model, the 1096 labelled images are sufficient for a segmentation of leaves and heads. For any organ, including stems, the increase in model performance levelled out when more than 60% (ca. 600 images) of the totally available data were used for training. This indicates that either a massively larger amount of data or a different labelling or training strategy might be needed for substantial further improvement. Model performance was reduced when performing a region-specific data split.

Among the two evaluated models, we chose Segformer. It achieved good segmentation for leaves, heads and background (IoU 80%), but the segmentation of the stems underperformed (IoU = 44.92%). The low performance of the stem segmentation may be attributed to various reasons: stems are thin, partially occluded by leaves, and a limited proportion of total pixels compared to the other organs. During booting and spike emergence (i.e. the expansion of the growing spike within leaf sheaths), the cylindrical structure of the stem is lost and the foil-like structure of the sheaths becomes obvious. This may lead to low model performance during this stage, particularly for stems. Furthermore, annotation difficulties related to distinguishing stems from some soil residues and senescent or rolled leaves may produce label noise. Improving stem segmentation may require exploring loss functions associated with class imbalance or incorporating massively more images with annotated stem masks.

These findings highlight a critical requirement: current models for segmenting organs in plants are highly dependent on extensive, high-quality annotations to achieve strong performance. To address this, research should prioritise the development of annotation-efficient solutions that maintain high performance with fewer labelled samples. Promising directions include leveraging self-supervised learning, which utilises unlabelled data and semi-supervised or active learning, which strategically selects a minimal number of samples for annotation while maximising model learning.

Without being quantitative, visual observation of the segmented image confirmed that wheat plants were segmented independently of their colour while weeds or plant residues were classified as background. This is a great step forward. Earlier models segmented plant tissue to a large degree based on colour. For example, in the case of Eschikon wheat segmentation training (EWS) [18], necrotic leaves were no longer detected as part of the canopy. This led to a decrease in canopy cover after winter due to necrotic leaves that suffered freezing damage [18]. Although the VegAnn model [9] included necrotic parts of plants in its segmentation while excluding crop residues, it was trained on a large number of different species and did not exclude weeds. [34]

### 4.5 Value of the full GWFSS dataset without labels

Self-supervised learning (SSL) methods can leverage a large amount of unlabelled image data as a pre-training procedure to better initialize or condition a deep learning model for a subsequent downstream analysis task. Ogidi et al. [34] found that a diverse source dataset in the same domain or similar as the target dataset combined with SSL can maximize performance in downstream plant phenotyping tasks. Our large, unlabelled dataset of 52,078 RGB images is meant as a training set for SSL methods. The idea of this dataset is to provide it as set for SSL while using the labelled data for validation and testing. The dataset can be also used for further labelling without the need to collect the data. It may be used for further segmentation tasks such as spikelet segmentations or other canopy features.

Beyond organ segmentation, the extensive dataset documented here provides a dynamic platform to develop predictive models that can capture temporal and spatial variability across multiple years and environments. By integrating environmental data with image-derived traits and machine learning methods, such as random forest regression or XGBoost, which can handle a vast array of predictor variables, breeders can target more complex traits such as radiation use efficiency, harvest index and yield. More fundamental underlying traits can be capable of better account for genotype-by-environment interactions and permit breeding programs to optimise their pipelines on a global scale.

### 4.6 Conclusion and outlook

Segmentation models will reduce the subjectivity of field observations by leveraging the generation of large and consistent datasets. Organ segmentation will enable the extraction of a range of additional traits from complex canopies. Such information is needed to advance our understanding of the interaction of *Genotypes* with the *Environment* they grow in and the *Management* practices they receive (often abbreviated as GxExM). To achieve robustness, the training data for the segmentation models needs to be large and diverse. Models derived from the GWFSS dataset will likely outperform many models derived from labelling in single experiments. As a further step, datasets from other small-grain cereals, such as barley, could be considered to enhance the training data.

While organ proportions will be directly available from the GWFSS-derived segmentation models, other traits will need to be developed and calibrated through secondary processing. For example, sensor fusion may allow one to integrate organ information derived from point sensors or sensors with lower resolution. Moreover, organ information may complement canopy-level traits derived from remote sensing.

Many of our images were derived from fully automated platforms, and these installations are often prototypes. With the advancement of agricultural robots, such platforms will become affordable for a greater community in the near future. But also smartphones or specifically designed handheld devices, will bring image-based phenotyping to a greater community. This will leverage new possibilities for common research projects in science and citizen science communities.

## Glossary

**Table 6:** Glossary of relevant terms used.

| Term | Abbreviation | Description | Reference |
| --- | --- | --- | --- |
| Labels |  | GWFSS context: polygons used to generate masks for the different organs. |  |
| Tags |  | GWFSS context: keywords added to an image, such as the developmental stage displayed. |  |
| Tiles |  | GWFSS context: subsamples from the original image that are used for labelling and training. Standard tiles are squares, often with 256 x 256 pixels. |  |
| Uniform Manifold Approximation and Projection | UMAP | Dimension reduction technique that can be used for latent feature visualisation | McInnes, L, Healy, J, UMAP: Uniform Manifold Approximation and Projection for Dimension Reduction, ArXiv e-prints 1802.03426, 2018 |
| Intersection over Union | IoU | IoU is a metric of segmentation performance of the area manually labelled (A) vs. the area detected by the model (B). The intersection size between the two is divided by their union size.<br>$\frac{\text{intersection}}{\text{union}}$ | https://en.wikipedia.org/wiki/Jaccard_index |
| Proximal sensing |  | Sensing from proximity. Sensors are typically operated hand-held or mounted on poles, gantries, or ground-based vehicles. |  |
| Imaging setup |  | The combination of vector (carrier systems), sensor and lens (i.e. RGB camera-lens combination) as well as the working distance and camera angle defining the field of view. | https://en.wikipedia.org/wiki/Field_of_view<br>vor definition of the field of view. |
| Phenology |  | The study of periodic events in biological life cycles and how these are influenced by seasonal and interannual variations in climate, as well as habitat factors (such as elevation) | https://www.merriam-webster.com/dictionary/phenology |
| Canopy |  | Branches (stems), leaves, and spikes (inflorescences) of a population of plants growing on a piece of land. | Adapted from different sources <sup>6</sup> |
| Crop ontology | CO | Provides descriptions of agronomic, morphological, physiological, quality, and stress traits along with their definitions and relationships. | https://cropontology.org |
| Leaf area index | LAI | Trait, which characterise plant canopies, are defined as leaf green area per unit of surface area | CO.321:0000184 |
| Initiation of booting | Boot | Phenological period prior to spike emergence where the flag leaf is fully developed | CO.321:0000191 |
| Anthesis | Ant | Phenological period when pollination occurs in wheat | CO.321:0000121 |
| Heading | Hd | Phenological period from the time of emergence of the spike tip from the flag leaf until the spike has fully emerged | CO.321:0000007 |
| Maturity | Mat | Phenological stage when wheat stops remobilising assimilates to the spike | CO.321:0000022 |

## Acknowledgments

We thank all field staff who ran the experiments and collected the images used in this study.

## Author Contributions

The GWFSS consortium consisted of different working groups that focused on conceptualisation and steering, data collection and data supply, labelling, data curation, training the base model, and writing (Table 7). The display of the author’s contribution was inspired by ^7^.

**Table 7:**
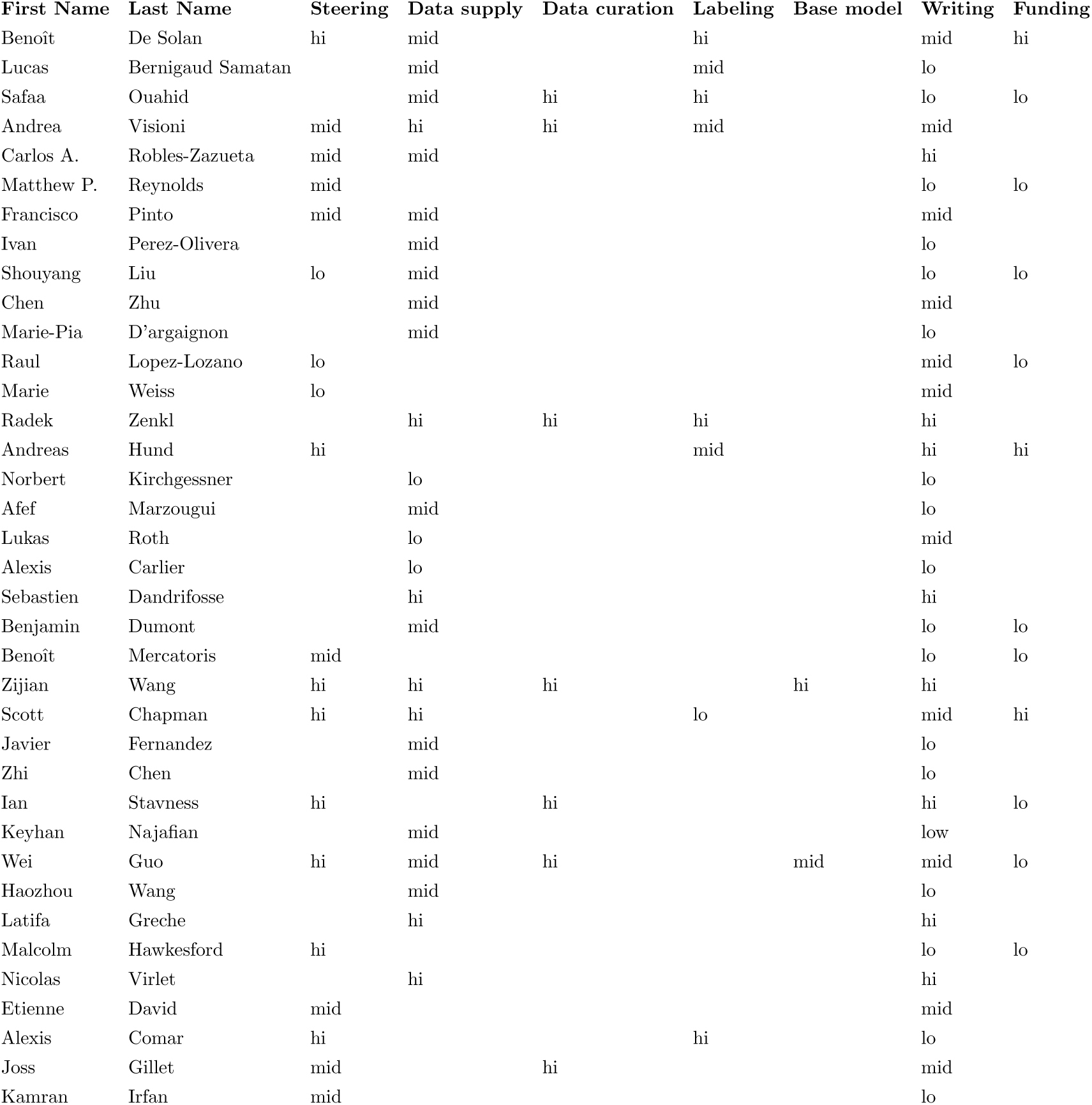
Author contribution.

## Funding

**Global wheat** was directly supported by Analytics for the Australian Grains Industry (AAGI). AAGI (UOQ2301-010OPX) is a Strategic Partnership between the Grains Research and Development Corporation (GRDC), Curtin University, The University of Queensland and the University of Adelaide. Other project and associate partners also support the initiative; Arvalis, France; Phenet (European Commission [96]); EMPHASIS-GO (European Commission [97]); Delley Seeds and Plants Ltd, Switzerland; Deutsche Saatveredelung AG, Germany; ASUR plant breeding, France; Plant Phenomics journal, China.

### Funding of individual projects of partners

**CIMMYT**: The International Wheat Yield Partnership (IWYP); the Heat and Drought Wheat Improvement Consortium (HeDWIC); the Accelerating Genetic Gains in Maize and Wheat (AGG); Modernizacion Sustentable de la Agricultura Tradicional (MasAgro) an initiative from the Secretaria de Agricultura y Desarrollo Rural (SADER), Mexico; Foundation for Food and Agricultural Research (FFAR). **ETHZ**: Swiss National Science Foundation (SNSF) **INRAe**: FFAST (French National Research Agency, ANR project number ANR-21-CE45-0037). **RRes**: Biotechnology and Biological Sciences Research Council (BBSRC) of the UK as part of the project Delivering Sustainable Wheat (BB/X011003/1). **Ulìege**: National Fund of Belgium F.R.S-FNRS (FRIA grant), Agriculture, Natural Resources and Environment Research Direction of the Public Service of Wallonia (project D31-1385 PHENWHEAT). **UQ**: INVITA - A technology and analytics platform for improving variety selection, GRDC UOQ2003-011RTX and GRDC UOQ2002-08RTX High-throughput feature extraction from imagery to map spatial variability. **USask**: the Natural Sciences and Engineering Research Council of Canada (NSERC) and the Canada First Research Excellence Fund (CFREF). **UTokyo**: the Japan Science and Technology Agency (JST) AIP Acceleration Research (JPMJCR21U3), the Sarabetsu Village “Endowed Chair for Field Phenomics” project in Hokkaido, Japan.

## Data Availability

The full dataset (GWFSS v1.0 full) including the 1,096 ground-truth labelled images (GWFSS v1.0 labelled), the descriptions of the datasets (GWFSS v1.0 subsets.csv) and imaging setups (GWFSS v1.0 imaging setups.csv) is available in the ETH research collection (http://dx.doi.org/10.3929/ethz-b-000734546) and should be referred to by citing this publication. The 1,096 labelled images will be released on July 15, 2025. To facilitate access, the labelled data and the benchmark model will also be available at (https://huggingface.co/datasets/GlobalWheat/GWFSS_v1.0). Further subsets of the data are available as part of a competition held in the framework of the Seventh International Workshop on Machine Learning for Cyber-Agricultural Systems (MLCAS2025). These sets are listed in Annex 1.1 but are not part of the publication. Links to these datasets can be found at: https://www.global-wheat.com/gwfss.html.

## 1 Annex

### 1.1 Data Challenge

By sharing the *full* GWFSS dataset and *tiny* set with the annotations, the objective is to enable the computer vision community to design wheat-related segmentation models, including instance segmentation. In particular, the full data set should encourage the development of tools that account for the complex structure of wheat, which can assist in extracting small features in dense vegetation. Recognising the potential of interdisciplinary collaboration in plant phenomics, we propose a data challenge based on the GWFSS dataset. This challenge complements the previous GWHD project, which focused on object detection, by expanding the scope to include segmentation.

This challenge aims to bring together expertise from the phenomics and computer vision communities to explore innovative strategies that mitigate the dependency on large-scale annotations. As pixel-level labelling is both expensive and time-consuming, one of the key objectives of this competition is to explore the potential of leveraging vast unlabelled data to enhance model performance. In this challenge, participants are provided with 65,000 unlabelled images (512×512), cropped from 2,000 full-sized images (Fig. S1). Additionally, a set of 100 labelled images is provided, obtained through stratified sampling from all institutions except Arvalis, which serves as the test set. Mean IOU will be served as the evaluation metric for this competition. By fostering solutions that balance performance and annotation efficiency, the challenge seeks to advance segmentation research while addressing practical limitations. Details of the data challenge can be found via the global wheat website (https://www.global-wheat.com/gwfss.html).

**Figure S1:**
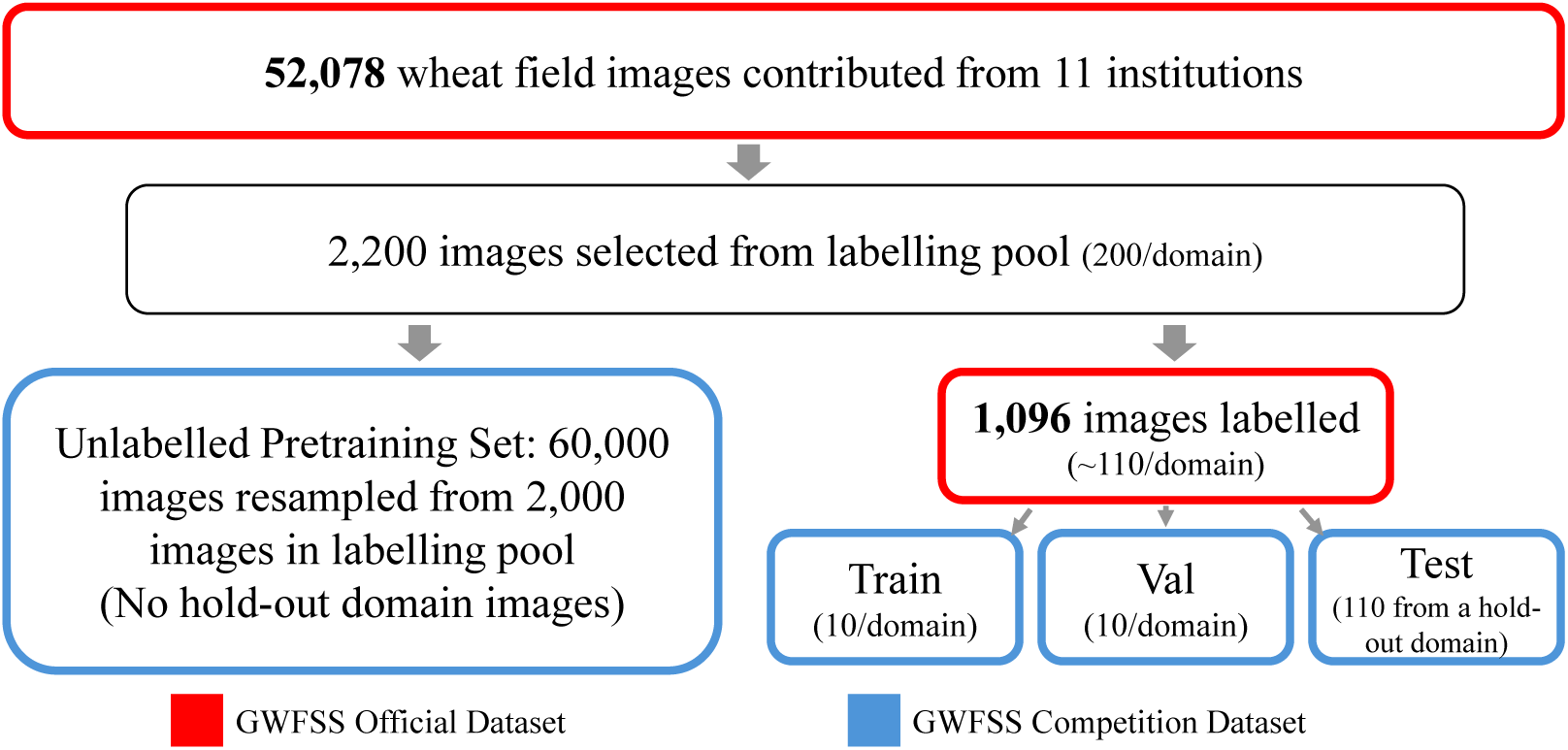
Workflow of dataset selection.

**Table S1:** Summary of all GWFSS variants.

| Dataset | Annotated | Split | Resolution | Number |
| --- | --- | --- | --- | --- |
| Full | Yes | - | $512 \times 512$ | 1096 |
| Full | No | - | Full Resolution (Tab.5) | 52,078 |
| Challenge | Yes | Train | $512 \times 512$ | 99 |
| Challenge | No | Train | $512 \times 512$ | 65,000 |
| Challenge | Yes | Val | $512 \times 512$ | 99 |
| Challenge | Yes | Test | $512 \times 512$ | 110 |

### 1.2 Evaluation of images containing weeds

**Figure S2:**
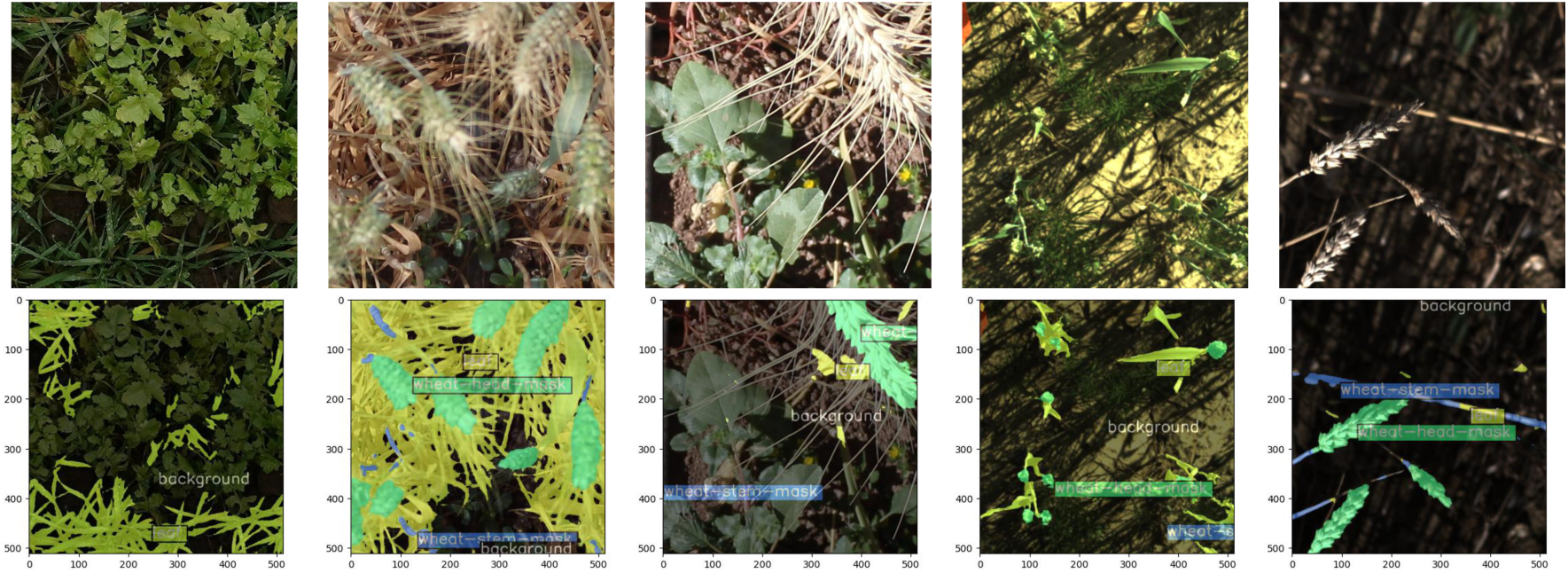
Visualisation of prediction results of all five images that contain weeds in the test set (Random Split, as described in Section 2.5.1)

### 1.3 Evaluation of images during the senescence phase

**Table S2:** Comparison of Mean Intersection over Union (mIoU) and mean accuracy (mAcc) metrics for Deeplabv3plus (R101) and Segformer (B1) across Random- and Region-based data splits.

|  | Later Stages |  | Full |  |
| --- | --- | --- | --- | --- |
|  | IoU | Acc | IoU | Acc |
| **Background** | 73.5 | 87.76 | 84.51 | 92.75 |
| **Head** | 85.21 | 90.65 | 82.85 | 90.14 |
| **Stem** | 43.35 | 51.72 | 44.92 | 53.89 |
| **Leaf** | 73.65 | 84.85 | 82.35 | 90.44 |
| **Average** | 68.92 | 78.74 | 73.66 | 81.81 |

**Figure S3:**
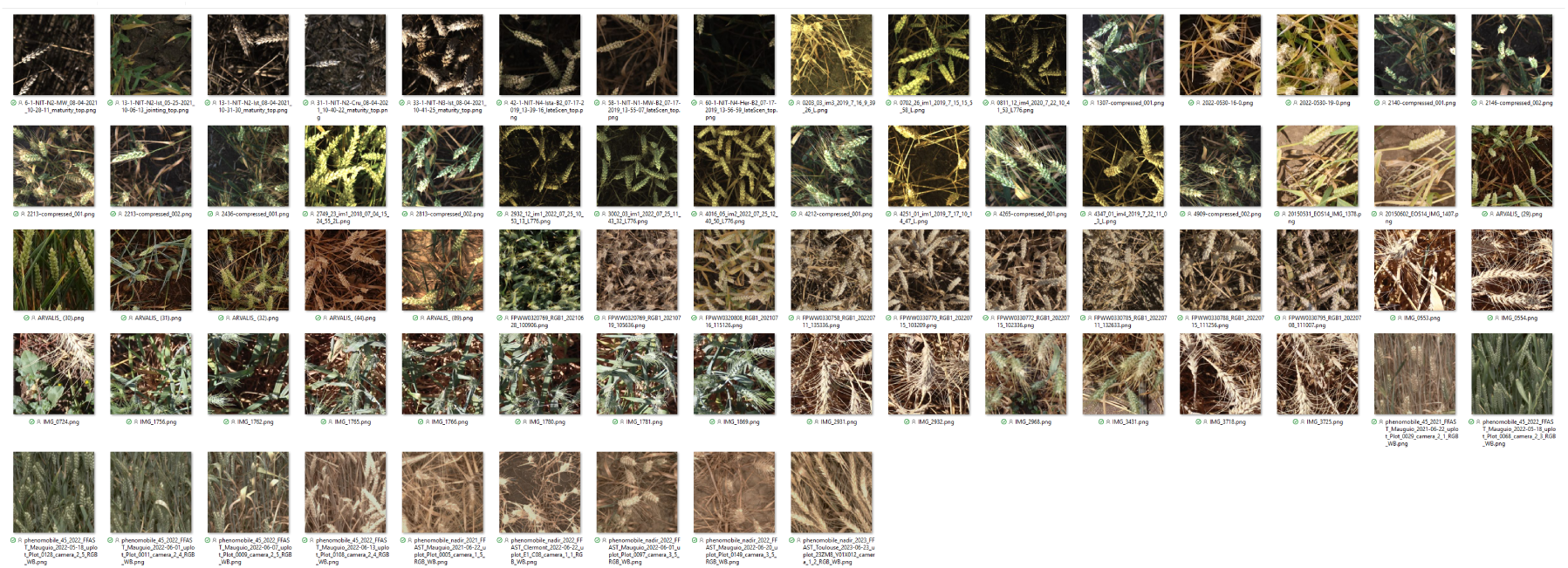
Images of the senescent set selected from the random split (Section 2.5.1) to evaluate model performance on canopies with chlorotic and necrotic tissue.

## Footnotes

1 https://darwin.v7labs.com

2 https://bioportal.bioontology.org/ontologies/BTO

3 https://www.ebi.ac.uk/ols4

4 https://cropontology.org/

5 https://bioportal.bioontology.org/ontologies/BTO

7 https://www.nature.com/nature-index/news/researchers-embracing-visual-tools-contribution-matrix-give-fair-credit-authors-scientific-papers

